# SCAR: Controlled mutations, ancient DNA damage, and fragmentation of fasta and fastq sequences

**DOI:** 10.64898/2026.09.24.754018

**Authors:** Mads Hartmann, Michael V Westbury

## Abstract

**Summary:** Controlled modification of sequencing data is important for reproducible benchmarking, particularly when evaluating analyses affected by read fragmentation, divergent reference genomes and ancient DNA damage. SCAR introduces controlled mutations, fragmentation and position-specific damage into FASTA and FASTQ data. All features can be used separately or in combination, and empirical fragment-length and mismatch profiles can be supplied to closely reproduce the properties of targeted datasets. Validation analyses showed the successful implementation of the requested sequence changes, and their impact on read mapping and heterozygosity analyses.

**Availability and Implementation:** SCAR is implemented in C++ and the latest code is available at https://github.com/Madshartmann1/SCAR. The version described in this article is archived at Zenodo at https://doi.org/10.5281/zenodo.22789736, with validation material available at https://doi.org/10.5281/zenodo.21915766.

## 1 Introduction

Simulations are valuable for developing and evaluating bioinformatic methods. In contrast to empirical data, simulated datasets provide a known ground truth and allow individual parameters and analytical choices to be tested under controlled and reproducible conditions. Useful benchmarking therefore requires simulations that are both configurable and representative of real sequencing data. This is particularly relevant for read mapping, variant calling, reference bias evaluation and analyses involving ancient DNA, where sequencing errors, biological variation, fragmentation and post-mortem damage can bias genomic inference [15].

Currently, there are several tools to simulate sequencing reads. For example, ART [6], wgsim [9], and Gargammel [12]. These tools generate synthetic sequencing reads from reference sequences, with ART modelling platform specific sequencing errors and Gargammel additionally simulating characteristics of ancient DNA. However, they primarily use consensus reference sequences and have limited capability for introducing variation into empirical sequencing reads and genomic datasets.

A tool that could instead modify empirical or previously simulated FASTQ reads directly would allow defined sequence changes to be introduced against the existing background of biological variation and sequencing errors, retaining the original dataset as a baseline for comparison. Mapping behaviour could then be compared before and after fragmentation, ancient DNA damage or introduced variation, and the resulting effects on reference bias and on analyses, such as heterozygosity estimation, quantified against that baseline. Such estimates are central to inferences about inbreeding (runs of homozygosity), genetic load (masked load) and demographic history (e.g. PSMC[10]), so understanding how data quality affects them is especially important for assessing genomic health and in the field of conservation genomics.

An approach of this kind should be applicable to reference sequences as well as to existing read data, allowing mutations, fragmentation and ancient DNA damage to be introduced independently or in combination.

Here, we present SCAR (Sequences + Controlled mutations + Ancient damage + fRagmentation), a modular program for introducing controlled mutations, sequence fragmentation and position dependent ancient DNA damage into FASTA and FASTQ files. SCAR supports multiple models for mutations, including fixed or rate based substitutions, transition-transversion biases, and user defined mutation spectrums, and records introduced mutations in a separate SNP receipt. Ancient DNA damage can be applied with predefined profiles or via empirical mapDamage [7] output files, while fragmentation can follow fixed, parametric or empirical length distributions.

## 2 Features and Methods

SCAR modifies existing FASTA or FASTQ sequences through three modular operations: fragmentation, mutation and ancient DNA damage (Figure 1). Each operation may be used independently or in combination. When combined, they are applied in a fixed order— fragmentation, mutation then ancient DNA damage—ensuring that mutations and terminal damage are introduced relative to the final fragments.

**Figure 1.**
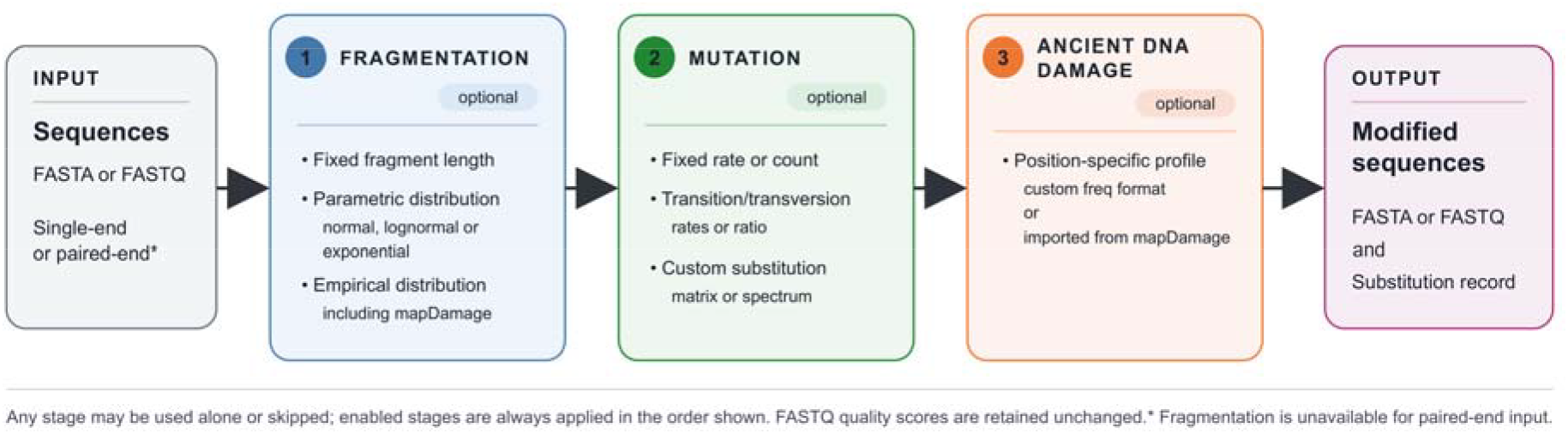
Flowchart for SCAR’s main functions. An input of FASTA or FASTQ sequences go through one or more of the methods; fragmentation, mutation and/or ancient DNA damage. These can be independent or combined, but will happen in the order shown here. The output is accompanied but a record of all introduced substitutions, in .snp format. Fragmentation is unavailable for paired-end inputs.

### 2.1 Features

#### 2.1.1 Fragmentation

SCAR supports fragmentation of long sequences using either a fixed fragment length or user defined fragment length distributions. Supported distributions include exponential, normal and lognormal models, as well as empirical distributions. Empirical input may be supplied as an unweighted list of observed lengths, a weighted length table or a mapDamage length distribution file. During processing, fragment lengths are sampled successively until the input sequence is consumed, with terminal remainders retained only when they exceed a user defined minimum length. To preserve read pairing for mapping, fragmentation is restricted to single-end processing.

#### 2.1.2 Mutation

SCAR supports several modes for nucleotide substitutions. Mutations can be introduced using a uniform per base rate or as an exact number across the dataset. Transition and transversion frequencies can be specified independently or derived from an overall mutation rate and a user defined transition-to-transversion ratio. For more detailed simulations, users can provide a custom mutation matrix, defining the rate of each of the 12 possible nucleotide substitutions or a custom mutation spectrum can define the relative frequencies.

For each selected position, the original nucleotide is replaced according to the chosen mutation model. Positions containing ambiguous nucleotides are not modified. Introduced substitutions are recorded in a tab separated SNP receipt containing the sequence identifier, 1 based position, original nucleotide and derived nucleotide. FASTQ quality scores are retained without modification.

#### 2.1.3 Ancient DNA damage

SCAR can simulate position dependent ancient DNA damage patterns characteristic of both double-stranded and single-stranded libraries. These include C→T substitutions enriched towards the 5′ end and G→A substitutions towards the 3′ end in double-stranded libraries, and C→T substitutions enriched towards both ends in single-stranded libraries. Substitution probabilities can be specified for individual positions from each fragment terminus using either user defined profiles or empirical profiles derived from mapDamage.

Damage profiles can be supplied directly or imported from a mapDamage output directory, with the format detected and converted automatically. For each applicable nucleotide, damage is introduced according to the position specific probability in the supplied profile. An optional background substitution rate can additionally be applied to positions not modified by the damage profile.

### 2.2 Software implementation

SCAR is implemented in C++ and uses a producer worker structure to divide sequence processing across CPU cores. Single-end FASTQ files can be processed in streaming mode with limited memory use, while paired-end mode processes synchronised read pairs and retains their original order. Random changes are controlled by a user defined seed, with per read seeds used to keep results reproducible across thread counts. Output compression follows the input format by default, and introduced substitutions are recorded in separate SNP files. The modular structure separates input handling, fragmentation, mutation and damage, making it straightforward to add further processing modes if needed.

### 2.3 Validation setup

SCAR was evaluated using three publicly available Lion *(Panthera leo)* datasets from SRA, their accession numbers are ERR13719741, ERR13719744 [3] and SRR27226083 [2].

We processed the reads with SCAR using mutation, fragmentation and ancient DNA damage, both separately and in selected combinations. For mutation only conditions, R1 and R2 were retained as paired-end reads and processed separately. Because SCAR applies fragmentation to single-end input, read pairs used in fragmentation containing conditions were matched by normalised read name and converted into single sequences by concatenating R1 with the reverse complement of R2; the R2 quality string was reversed accordingly. This procedure concatenated the complete mate sequences rather than merging them according to sequence overlap. Modified and unmodified reads were subsequently mapped against the lion reference genome P.leo_Ple1_pat1.1 (GCF_018350215.1; [1]) using PlainMap (version 0.1.0) [13] with default parameters.

Validation assessed whether SCAR introduced the requested sequence properties while preserving valid FASTQ structure, read identifiers and quality scores. Empirical fragment length distributions and damage profiles were derived from a double-stranded library from an El Olivar camelid (sample 14B; [11]) and a single-stranded library from a cave hyena (sample Ccsp015; [14]).

Fragmentation was evaluated by comparing the specified and resulting fragment length distributions. Ancient DNA damage was evaluated from position specific C→T and G→A misincorporation frequencies estimated with mapDamage [7](version 2.2.3). Mutation modes were evaluated from the recorded substitution counts and spectra. We estimated genome wide heterozygosity using allele frequencies (−doSaf 1) in ANGSD [8] (version 0.941-26-g6b5d906), taking genotype likelihoods into account using the GATK model (−GL 2). We retained uniquely mapped reads (−uniqueOnly 1), excluded reads flagged as bad (−remove_bads 1), required a minimum mapping quality of 30 (−minMapQ 30) and minimum base quality of 20 (−minQ 20), adjusted quality scores around indels (−baq 1), excluded transitions (−noTrans 1), and removed sites with a total depth below 5X. To ensure comparability between conditions with differing coverage, all sample condition combinations were downsampled to an expected mean depth of 14.56X using ANGSD, corresponding to the lowest observed depth. Heterozygosity was computed from the resulting site allele frequency likelihoods using realSFS from the ANGSD toolsuite, analysing batches of up to 5,000,000 covered sites (−nSites). For each sample and condition, heterozygosity was calculated as the heterozygous site frequency spectrum bin divided by the sum of all three bins across batches. For the ancient DNA condition heterozygosity estimates, only the data with double stranded simulated damage was used.

## 3. Results and discussion

In our validation dataset, SCAR applied the requested sequence modifications across all three modes. Mutation introduced substitutions were in accordance to the specified rates and spectra, with introduced substitutions correctly recorded in the corresponding SNP receipts. Fragmentation shifted mapped read length distributions from median lengths of 150-151 bp in the original datasets to 47-48 bp, closely matching SCAR’s input fragment-length distribution after mapping (Figure 2a). Ancient DNA damage generated the expected position dependent misincorporation patterns for both library preparation types: C→T substitutions increasing towards the 5′ end, with complementary G→A substitutions towards the 3′ end for the double-stranded profile, and C→T substitutions increasing towards both fragment termini for the single-stranded profile. These patterns were retained after mapping and were recovered with mapDamage, closely matching the input profiles at both fragment termini for each library type (Figure 2b-d). For the double-stranded profile, 3′ C→T misincorporation is not an expected damage signal, so misincorporations there likely reflect background noise and variants relative to the reference genome, rather than true damage, consistent with its comparatively narrow y-axis scale.

**Figure 2.**
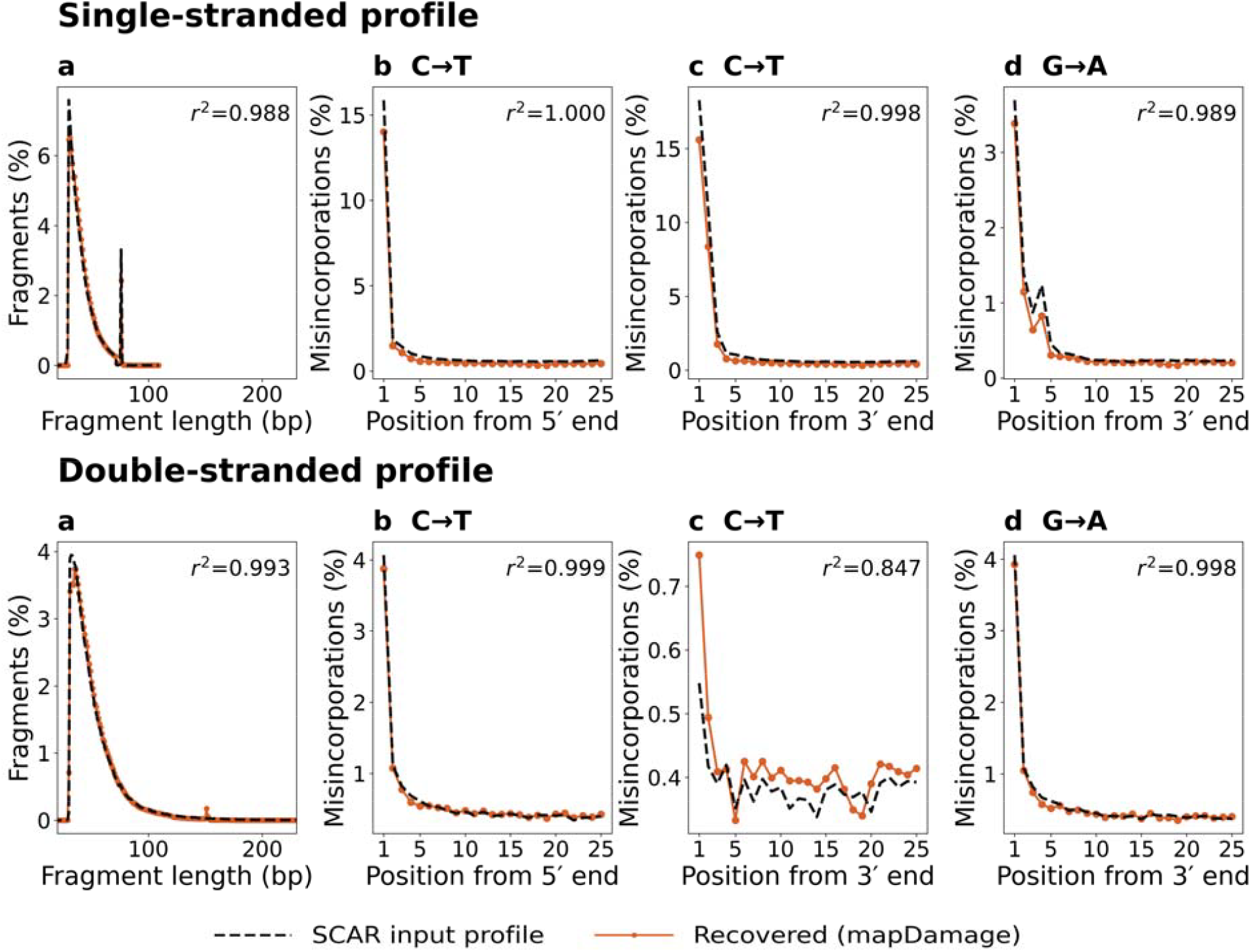
Recovery of SCAR’s input fragmentation and ancient damage profiles after mapping, for single-stranded (top row) and double-stranded (bottom row) damage profiles, using accession ERR13719744. Dashed black lines show the input profile supplied to SCAR; coloured lines show the profile recovered from mapDamage after mapping. **a**, fragment-length distribution. **b**, 5′ C T misincorporation frequency. **c**, 3′ C T misincorporation frequency. **d**, 3′ G A misincorporation frequency. Pearson between the input and recovered profiles is given in each panel.

Mutation alone had little effect on mapping depth or breadth of coverage, whereas conditions including fragmentation reduced mapping rates and coverage across datasets (Table 1). Mean depth decreased in two datasets but increased in ERR13719741. This depth reduction is expected as SCAR discards the terminal remainder of a read during fragmentation whenever it falls below the minimum fragment length, as described earlier. As a result a fraction of otherwise usable bases are lost at the fragmentation step itself, rather than being retained as a shorter mapped fragment. We further note that fragmentation reduced the duplication rate from 38.12% to 16.81% in ERR13719741. This suggests that identical reads fragmented at different positions may subsequently have been treated as unique, indicating that duplicate removal before fragmentation may be preferable for highly duplicated datasets.

**Table 1.** Mapping rate and mean depth across the four SCAR processing stages for the three validation accessions. aDNA denotes the combined fragmentation and ancient damage simulation condition.

| Accession | Raw | Mutated | Fragmented | aDNA |
| --- | --- | --- | --- | --- |
| ERR13719744 | 93.1% (17.2 ) | 92.8% (17.2 ) | 76.4% (15.9 ) | 76.0% (15.8 ) |
| SRR27226083 | 92.7% (34.7 ) | 92.3% (34.7 ) | 73.9% (25.3 ) | 73.5% (25.2 ) |
| ERR13719741 | 94.6% (14.5 ) | 94.5% (14.5 ) | 78.3% (17.3 ) | 77.9% (17.2 ) |

Following depth normalisation and transition filtering, mutation alone caused only modest increases in estimated heterozygosity, whereas conditions including fragmentation or ancient DNA damage produced substantially larger reductions (Table 2). These results show that SCAR-induced sequence modifications can measurably affect downstream mapping and heterozygosity estimates.

**Table 2.** Estimated heterozygosity (X 10^−4^) across the four SCAR processing stages for the three validation accessions, with percent change relative to the untreated (OG) value shown in parentheses.

| Accession | Raw | Mutated | Fragmented | aDNA |
| --- | --- | --- | --- | --- |
| ERR13719744 | 1.318 | 1.340 (+1.7%) | 1.084 (−17.7%) | 1.042 (−21.0%) |
| SRR27226083 | 6.414 | 6.910 (+7.7%) | 2.940 (−54.2%) | 2.747 (−57.2%) |
| ERR13719741 | 1.443 | 1.498 (+3.9%) | 1.279 (−11.3%) | 1.229 (−14.8%) |

The heterozygosity comparisons illustrate why controlled simulations are useful for addressing this kind of question. All conditions were compared at the same mean depth and with transitions excluded, so the differences in Table 2 cannot be explained by differences in mean sequencing depth alone, nor by damage derived bases being scored as heterozygous sites. Even so, fragmentation and ancient DNA damage reduced estimated heterozygosity by up to 54% and 57%, respectively, whereas substitutions introduced directly into the reads changed the estimates by only a few percent. Fragmentation and damage therefore affected the heterozygosity estimates more than the introduced sequence divergence. This is most likely because shorter and damaged fragments change which regions remain mappable and which sites pass the analysis filters. The size of the effect also differed markedly between the three lion individuals, indicating that the biases may be dataset specific. Given that heterozygosity is used to infer inbreeding, genetic load and demographic history, differences of this size may affect biological conclusions when datasets differ in quality/ preservation or fragment length.

Because SCAR modifies the reads themselves, the unmodified dataset can be used as a direct comparison when testing the effect of these changes for a specific sample, reference genome and/or analysis pipeline. A well preserved modern dataset can, for example, be degraded using the fragment length and damage profiles of an ancient sample, supplied directly as mapDamage output, to estimate how much of the difference between the datasets can be explained by data quality alone. The same principle can be used to investigate the effect of reference divergence by introducing a defined number of substitutions into the reference genome and comparing the proportion of reads recovered when mapping to the modified and original references. SCAR therefore complements existing read simulators rather than replacing them. Sequencing characteristics can be retained from empirical data or generated using existing tools, while mutations, fragmentation and post-mortem damage are added afterwards in a controlled and reproducible order, either separately or in combination.

## 4. Data Availability Statement

SCAR source code and documentation are available at https://github.com/Madshartmann1/SCAR. The version of SCAR used in this study is archived at Zenodo [4]. Validation settings, commands, scripts and results are available separately at Zenodo [5]. The sequencing datasets are available from the SRA under accessions ERR13719741, ERR13719744 and SRR27226083.

